# Semaglutide Improves Myocardial Perfusion and Performance in a Large Animal Model of Coronary Artery Disease

**DOI:** 10.1101/2024.08.15.608191

**Authors:** Christopher R. Stone, Dwight D. Harris, Mark Broadwin, Meghamsh Kanuparthy, Ju-Woo Nho, Keertana Yalamanchili, Jad Hamze, M. Ruhul Abid, Frank W. Sellke

## Abstract

**Objective:** Coronary artery disease (CAD) is the leading cause of death worldwide. It imposes an enormous symptomatic burden on patients, leaving many with residual disease despite optimal procedural therapy, and up to 1/3 with debilitating angina amenable neither to procedures, nor to current pharmacologic options. Semaglutide, a glucagon-like peptide 1 agonist originally approved for management of diabetes, has garnered substantial attention for its capacity to attenuate cardiovascular risk. Although subgroup analyses in patients indicate promise, studies explicitly designed to isolate the impact of semaglutide on the sequelae of CAD, independently of comorbid diabetes or obesity, are lacking.

**Approach and Results:** Yorkshire swine (n=17) underwent placement of an ameroid constrictor around the left circumflex coronary artery to induce CAD. Oral semaglutide was initiated postoperatively at 1.5 mg and scaled up in 2 weeks to 3 mg in treatment animals (SEM, n=8) for a total of 5 weeks, while control animals (CON, n=9) received no drug. All then underwent myocardial harvest with acquisition of perfusion and functional data using microsphere injection and pressure-volume loop catheterization. Immunoblotting, immunohistochemistry, and immunofluorescence were performed on the most ischemic myocardial segments for mechanistic elucidation. SEM animals exhibited improved left ventricular ejection fraction, both at rest and during rapid myocardial pacing (both p<0.03), accompanied by increased perfusion to the most ischemic myocardial region at rest and during rapid pacing (both p<0.03); reduced perivascular and interstitial fibrosis (both p <0.03); and apoptosis (p=0.008). These changes were associated with increased activation of the endothelial-protective AMPK pathway (p=0.005), coupled with downstream increases in endothelial nitric oxide synthase (p=0.014).

**Conclusion:** This study is the first to reveal the capacity of oral semaglutide to augment cardiac function in the chronically ischemic heart in a highly translational large animal model, likely through AMPK-mediated improvement in endothelial function and perfusion to the ischemic myocardium.

## Introduction

Coronary artery disease (CAD), when considered in conjunction with consequent cardiac infarction and failure, is unique in both ubiquity and lethality. It is not only the current leading cause of death worldwide but is also, given its expansion to over 2 million new patients yearly over the past two decades, poised to persist as such indefinitely ^1–3^. Consensus regarding the gravity of these figures has precipitated commensurate expenditure on research and treatments aimed at their mitigation ^4^, but this has not, unfortunately, kept pace with the biological and societal impacts of CAD. This deficit of therapeutic provision relative to clinical need is instantiated in the rate of disease after employment of the established options, with persistence in 1/3^rd^ of patients who receive surgical revascularization, together with over ½ who undergo percutaneous stenting ^5^. Viewing this disparity in tandem with the sizeable number of patients with debilitating cardiogenic pain who are not candidates for the standard procedural regimen, a mandate to develop effective adjunctive approaches clearly emerges.

The promise conferred to fulfill this role by semaglutide, popularly designated by its trademarks *Ozempic, Wegovy, and Rybelsus*, is a function of several features of its biology that have been described in recent years. First characterized by Starling around the turn of the 20^th^ century, the incretin hormone GLP-1 from which semaglutide descends was originally investigated for its ability to augment glucose-dependent insulin secretion. Although initially discovered to act on targets in the pancreas and brain, receptors were subsequently found in the heart in both animals and humans, with patients exhibiting levels comparable to those in the pancreas in all cardiac chambers ^6,7^. It is thus not surprising that, in addition to its well-characterized glucose-lowering and weight-controlling effects, many favorable cardiovascular sequelae have been noted after GLP-1 agonist administration, including reduced blood pressure, lowering of lipid levels and inflammation, and, beginning in 2016 with the LEADER trial, reduction in cardiovascular risk ^8^. A meta-analysis of the results of eight randomized controlled trials examining the use of GLP-1 agonists in the setting of diabetes concluded that these agents reduced major adverse cardiac events and all-cause mortality in this population ^9^. These findings were recently replicated in an additional randomized trial of patients with obesity but without diabetes ^10^. There have been many studies designed to account for these effects mechanistically that collectively constitute a thorough account of the multitude of cardiovascular and metabolic targets favorably affected by the administration of GLP-1 agonist therapy. This appears to proceed both directly through action on the GLP-1 receptor, and as epiphenomena of changes in other cells ^11^: this latter category includes improved arteriolar eNOS activation, nitric oxide production, and vasodilation, as well increased coronary artery blood flow ^12,13^; reduced myocardial infarct size, apoptosis, and oxidative stress ^14^; and improved metabolic substrate handling in the ischemic heart ^15^.

Missing from this body of literature, however, even after inclusion of the comparatively sparse number of studies conducted in large animal models and in patients ^16–26^, is experimental efforts to address the consequences of GLP-1 agonist therapy on chronic ischemic heart disease directly. Currently available evidence is confined either to the acute context, in which GLP-1 agonists are administered after infarction is induced experimentally or experienced by patients; or to the comorbid context, in which diabetes, obesity, or often both, provide the rationale for GLP-1 agonist administration. While the data derived from these studies are central to the account of the cardiovascular benefits of GLP-1 agonists that has propelled them to their preeminent place in public estimation, scientific interest, and, most recently, FDA endorsement for the treatment of cardiovascular disease ^27^, they are also incomplete. A full description of the potential possessed by GLP-1 agonists for the treatment of CAD, in contrast, would require evidence gleaned from a protocol uncomplicated by the potentially confounding presence of additional disease states, and conducted over a timeline adequately prolonged to effectively model the chronicity of CAD. Because it conforms to these requirements, we drew on our established large animal model of isolated CAD to explore the effect of the GLP-1 agonist semaglutide on cardiac function, cardiovascular hemodynamics, coronary perfusion, and associated tissue- and molecular-level indicators of the myocardial response to chronic ischemic pathophysiology.

## Methods

### Animal Protocol Principles and Preparation (Week 1)

This experiment was designed, in acknowledgement of the importance of accounting for potential innate sex-based cardiac biological variability, to incorporate an equal number of male and female animals in both groups. Although our allocation satisfies this imperative, the sample sizes thus generated provide insufficient analytic power for the partitioning of animals into the sub-groups necessary to conduct sex-specific comparisons: using a standard deviation derived from our preliminary data and a b error of 0.10, a group size of 8 was derived. In contrast, the maximal group size available for sex-specific comparison in this experiment is 5. Given the countervailing ethical imperative to minimize the number of large animals used in surgical research, this represents an effort to balance conflicting considerations, ideally yielding an experiment that is both informative and responsibly conducted.

The protocol begins with the arrival of animals to our veterinary facility (Yorkshire swine, n=17, 8 female and 9 intact male, sourced from the Cummings School of Veterinary Medicine at Tufts University, Grafton, MA). Animals are aged 10 weeks on arrival, whereupon they are housed socially within limits imposed by their well-being, provided a standard porcine chow-based diet, and allotted a one-week period of acclimation to our facility prior to initiation of the experiment.

### Ameroid Constrictor Surgery (Week 2)

Aspirin at 10 mg/kg/day was administered to all swine for 1 day preoperatively and 5 days postoperatively to mitigate thrombotic risk and to maximize the consistency of ischemic territory size across animals ^28^. Intramuscular xylazine and telazol were provided for anesthetic induction, with isoflurane used to maintain anesthesia after endotracheal intubation was completed to minimize anesthesia-induced cardiovascular perturbation ^29^. After standard antiseptic preparation with alcohol and betadine, a left anterolateral thoracotomy was performed in the third thoracic interspace. A retractor was placed, and the pericardium overlying the left atrial appendage was thus brought into view; the pericardium was divided sharply and temporarily plicated to the chest wall to elevate the heart into the operative field. Upon exposure of the left atrial appendage, this structure was retracted laterally to reveal the left main coronary artery shortly distal to its origin at the aortic root. The bifurcation of the left main coronary artery into its left anterior descending and left circumflex (LCx) branches was identified, and the latter dissected free from the surrounding epicardial fat prior to its first obtuse marginal bifurcation. Heparin was administered at a dose of 80 IU/kg, and a silastic vessel loop was passed under the LCx and gently retracted for two minutes. While electrocardiographic monitoring during retraction confirmed occlusion through the production of ST segment elevation, a 5 mL solution of gold microspheres (BioPAL, Worcester, MA) was injected into the left atrial appendage over 30 seconds. 15 μm in diameter, these microspheres distribute to and become lodged in all coronary capillary beds, with the exception, due to the absence of native epicardial collaterals in the porcine heart ^30^, of those supplied by the LCx, as these are made inaccessible by occlusion produced by retraction of the vessel loop. This differential distribution subsequently provides the basis for use of a neutron activation assay to identify the tissue rendered most ischemic, as this tissue will hold the lowest quantity of gold ^31^. Following the two minutes allotted for occlusion, the vessel loop was removed and replaced with an ameroid constrictor (Research Instruments SW, Lebanon, OR) with an internal diameter chosen to approximate that of the isolated vessel. Nitroglycerin (1 mL) is applied topically to the manipulated vessel to mitigate spasm. Rotation of the ameroid to maintain its position despite the beating myocardium underneath was followed by closure of the pericardium with absorbable suture and then closure of the chest and skin in layers, completing the procedure.

### Semaglutide Therapy (Weeks 4-8)

The ameroid constrictor placed around the LCx on all animals is composed of an outer titanium ring that encircles an inner layer of hygroscopic casein. The inner casein layer closes gradually over the course of several weeks as it swells with pericardial fluid, replicating the gradual onset of CAD in patients ^32^. Initiation of semaglutide two weeks following ameroid placement therefore effectively replicates delivery of the drug following the establishment of CAD. Analogous to its use in patients, semaglutide was initiated in animals randomly allocated to receive this drug at half-dose at the beginning of the treatment period, and up-titrated to the full dose after toleration with minimal burden of the typical gastrointestinal side effects seen with this agent is established ^33^. Oral semaglutide was thus used, initially at 1.5 mg and subsequently at 3 mg for all animals, as none exhibited side effects. This dosing regimen represents half of the lowest dose given to patients, and was chosen in acknowledgement of both that the weight of animals in this experiment is approximately half that seen in the cardiovascular clinical trial published using this agent (40.5 kg vs. 90.9, respectively), and that this drug has never been administered in a comparable animal model ^33^. All animals were allocated to treatment (SEM, n=8) or control (CON, n=9) as an entire cohort prior to arrival and thus also prior to investigator knowledge of any relevant physical parameters, obviating the risk of bias in selection of animals for the experimental arm of the study. Both cohorts contained an approximately equal number of male and female animals (n=4 female in both, n=4 male in SEM, and n=5 male in CON).

### Myocardial Perfusion, Performance, and Resection (Week 9)

Following the 5^th^ week of either treatment with semaglutide or no drug, all animals underwent a terminal procedure for acquisition of cardiac functional parameters, perfusion studies, and that culminated in myocardial resection for sectioning and subsequent tissue- and molecular-level analysis. Perioperative sterile preparation and anesthesia were identical to the previous surgery, after which animals were positioned supine and subjected to median sternotomy for optimal exposure of the heart and great vessels. The pericardium was entered and adhesions to the heart were divided as needed until the bilateral atrial appendages were fully exposed. Pacemaker clamps emanating from a single-chamber temporary pacemaker (OSCOR, Palm Harbor, FL) were affixed to the appendages. Dissection continued within the pericardial sac to expose the left ventricular apex, and without to expose the inferior vena cava (IVC) at its entry into the right atrium; the latter was encircled with a silastic vessel loop to permit IVC occlusion.

With the heart safely dissected as needed for the remaining procedural steps, systemic heparinization with 80 IU/kg was given, and a femoral arterial cutdown and isolation was performed. A 6 French vascular cannula was inserted into the exposed artery and connected to a withdrawal pump (Harvard Apparatus, Holliston, MA). Blood was withdrawn using the pump and into a collection tube at 6.67 mL/min concurrently with injection of a 5 mL solution of isotope-labelled microspheres into the left atrial appendage, first at resting heart rate, and thereafter during pacing of the heart at 150 beats per minute; lutetium and/or europium was used at rest, and samarium and/or lanthanum was used during rapid pacing (all BioPAL, Worcester, MA). Identically for the gold microspheres used in the preceding procedure, post-operative employment of neutron activation permits quantification of microspheres within myocardial tissue. Placing this count in ratio with that of concurrently withdrawn blood then allows precise derivation of perfusion to the studied tissue, according to the following equation: (blood flow [mL/min]/tissue mass [g]) x (tissue microsphere quantity/blood microsphere quantity). Because this calculation is performed once for the isotope injected during resting heart rate, and then again for the isotope injected during rapid pacing, blood flow under resting conditions as well as during simulated stress is yielded using this method.

When microsphere injection and collection of isotopically infused blood was completed, the withdrawal pump was disconnected and replaced with a calibrated catheter (Miliar, Houston, TX) advanced through the femoral sheath for transduction of aortic pressure. An additional 6 French sheath was then inserted and secured into the left ventricular apical myocardium to accommodate a catheter calibrated to transduce pressure and volume through conversion from admittance (ADV500 Pressure-Volume Measurement System, Transonic, Ithaca, NY). This system generates pressure-volume loops in real time, providing the immediate intraoperative feedback needed to adjust catheter positioning for accurate recordings. Hemodynamic parameters were transduced by these means in triplicate during ventilatory pauses to attenuate the effect of respiratory variation on outputs under three conditions: resting heart rate, rapid pacing to 150 beats per minute, and following IVC occlusion. This range allows for the profiling of hemodynamics both across a spectrum of physiologic stress and, during inflow occlusion, in a load-independent fashion that permits calculation of intrinsic cardiac contractility ^34^.

After completion of hemodynamic characterization, catheters were withdrawn and hearts were resected for sectioning. The left ventricular myocardium was divided into apical and basal halves and then radially according to the position of the left anterior descending artery and the LCx. This technique generated ten left ventricular tissue sections, sub-sections of which were fixed in 10% formalin for subsequent histologic assays; snap frozen in liquid nitrogen for immunoblotting; or desiccated in preparation for neutron activation assays. This latter technique was performed first, as it allowed identification of the most ischemic section of left ventricular myocardium, which in turn served as the focus of subsequent assays.

### Metabolic and Physical Parameters

All animals were weighed prior to both surgical procedures. Length was also taken prior to the terminal procedure for derivation of body surface area, as this was needed to normalize hemodynamic parameters. Hematologic studies were obtained at the outset of the terminal procedure, with initial blood samples taken for baseline blood glucose, albumin, and serum lipids. After acquisition of baseline glucose, administration of dextrose (0.5 g/kg of 50% solution) allowed the performance of glucose tolerance testing at 30- and 60-minute timepoints thereafter.

### Immunofluorescence and Immunohistochemistry

Following delineation of the most ischemic segment of left ventricular myocardium using gold microsphere mapping, tissue from this segment, which had been previously fixed in formalin and then transferred after 24 h to 70% ethanol, was converted to blocks for application of immunofluorescent and immunohistochemical stains: Masson’s trichrome protocol was applied to both the interstitial and perivascular myocardium to delineate the degree of fibrotic change in these locations, and terminal deoxynucleotidyl transferase dUTP nick-end labeling (TUNEL) with fluorescein quantified comparative DNA cleavage as an indicator of apoptotic change. All staining was performed by unaffiliated technicians (iHisto, Salem, MA) blinded to experimental circumstances. Whole-slide quantification of immunofluorescent TUNEL was also performed by blinded and unaffiliated technicians using quantitative pathology software (Halo Indica, Albuquerque, NM), while immunohistochemical analysis of Masson’s trichrome was done by lab personnel using QuPath bioimage analysis software (Belfast, Northern Ireland, UK) accordingly to a previously established protocol ^35^. Briefly, interstitial fibrosis was calculated using a machine learning algorithm trained to specifically identify stained collagen fibers; this algorithm was then applied uniformly to all sides from both arms of the experiment to yield the mean fibrotic area per slide. The same algorithm was applied to homogeneously sized representative regions centered around 1-3 perpendicularly oriented blood vessels from all slides to calculate ratio of stained collagen to total perivascular area, whereby mean perivascular fibrosis per slide was generated.

### Immunoblotting

Tissue from the most ischemic left ventricular segment was also studied using immunoblotting for relative expression of several proteins of interest: phospho-Akt (p-Akt, Ser473), Akt, phospho-AMP-activated protein kinase-α (p-AMPK, Thr172), AMPK, phosphatidylinositol-3-kinase (PI3K, p110α), endothelial nitric oxide synthase (NOS), bcl-2- associated death promoter (BAD), cleaved caspase 3 (Asp175), apoptosis-inducing factor (AIF), and transforming growth factor-β (TFG-β, all Cell Signaling, Danvers, MA). Myocardial tissue was digested into lysates, the protein concentrations of which were quantified using radioimmunoprecipitation (BCA Protein Assay Kit, Thermo Fisher Scientific, Waltham, MA). Lysate concentration was then used to load 40 μg of total protein from each tissue section into wells of 4-12% Bis-Tris gels (Thermo Fisher Scientific, Waltham, MA). Gels were subjected to electrophoresis for protein separation and transferred to nitrocellulose membranes (Bio-Rad, Hercules, CA) overnight at 4°C, and then incubated over the subsequent night with primary polyclonal antibody solutions in 3-5% bovine serum albumin to the aforementioned targets at the same temperature. All membranes were additionally normalized for variation in loading by incubation with vinculin or glyceraldehyde-3-phosphate dehydrogenase (GAPDH, both Cell Signaling, Danvers, MA). After overnight incubation, primary antibody dilutions were washed from membranes, which were then incubated at room temperature for 1 hour with appropriate secondary antibody dilutions (anti-mouse or anti-rabbit, Cell Signaling, Danvers, MA). Following activation in chemiluminescent substrate (Thermo Fisher Scientific, Waltham, MA), marker signals were imaged in the ChemiDoc digital system (Bio-Rad, Hercules, CA), generating images for densitometric analysis using ImageJ (National Institutes of Health, Bethesda, MD). Repeat probing was performed as needed using Restore PLUS Western Blot Stripping Buffer (Thermo Fisher Scientific, Waltham, MA). Markers were placed in ratio to vinculin or GAPDH expression within the same well, and therefore reported as relative.

### Statistical Analysis

All computations were performed with Prism 10.1.0 statistical software (GraphPad Software, San Diego, CA). Prior to performance of comparisons, data distributions were assessed for normality using the Shapiro-Wilk method; normal distributions were compared using unpaired, two-tailed Student’s t tests, while the Mann-Whitney U test was employed for non-normal data. The presence of outlier was adjudicated using the ROUT and Grubbs methods using nonlinear regression and the extreme studentized deviate, respectively. A p-value threshold of 0.05 was pre-established to designate statistical significance but, given the small sample sizes and proportional power limitations entailed by large animal work, select differences between p=0.05 and p=0.10 were designated as trends in the context of additional corroborating data.

### Study Approval

The protocol described for this experiment was approved by and administered under the supervision of the Institutional Animal Care and Use Committee at Rhode Island Hospital. This governing body was tasked with ensuring that all experiments were conducted in conformity to the *Guide for the Care and Use of Laboratory Animals* that is promulgated by the National Research Council and was accorded inviolable authority to halt all scientific proceedings not adherent to this standard. There were no deviations from full compliance encountered during the conduction of this experiment.

### Materials and Data Availability

Any requests for additional elucidation of the means and materials by which this experiment was performed and by which its associated data was derived (e.g., raw pressure-volume loop tracings, microsphere counts used to derive perfusion data, and immunoblot densitometry values) will be honored upon request made to the corresponding author, who has access to all data and takes responsibility for the integrity and analysis thereof.

## Results

### Systemic and Metabolic Effects of Semaglutide

In alignment with the production by GLP-1 agonist therapy in patients of favorable modulations in serum lipids, glycemic control, and body composition (28, 29), we tracked physical and metabolic parameters in our experimental animals. At the outset of the experiment, animals that received semaglutide (SEM) weighed significantly less than control counterparts (CON, 18.2 kg vs. 25.5, respectively p=0.042); at the end of the protocol, SEM animals continued to weigh less than the CON mean, but the difference lost statistical significance (37.3 vs. 44.4 kg, respectively, p=0.103). There was no difference in weight change over the course of the protocol (p=0.398). In contrast, there was a significant difference in body surface area favoring treated animals at the conclusion of the protocol (0.87 vs. 1.0 m^2^, p=0.029). Glucose tolerance testing performed at the harvest procedure revealed no difference at baseline or at 30 minutes, but a significant improvement at 60 minutes (189 mg/dL vs. 236 mg/dL, p=0.013). Serum triglycerides were also significantly reduced in the SEM group (12 mg/dL vs. 19 mg/dL, p=0.023), while there was no difference in serum albumin between treatment groups (p=0.433). Table 1 represents these changes alongside their statistical interpretation.

**Table 1.**
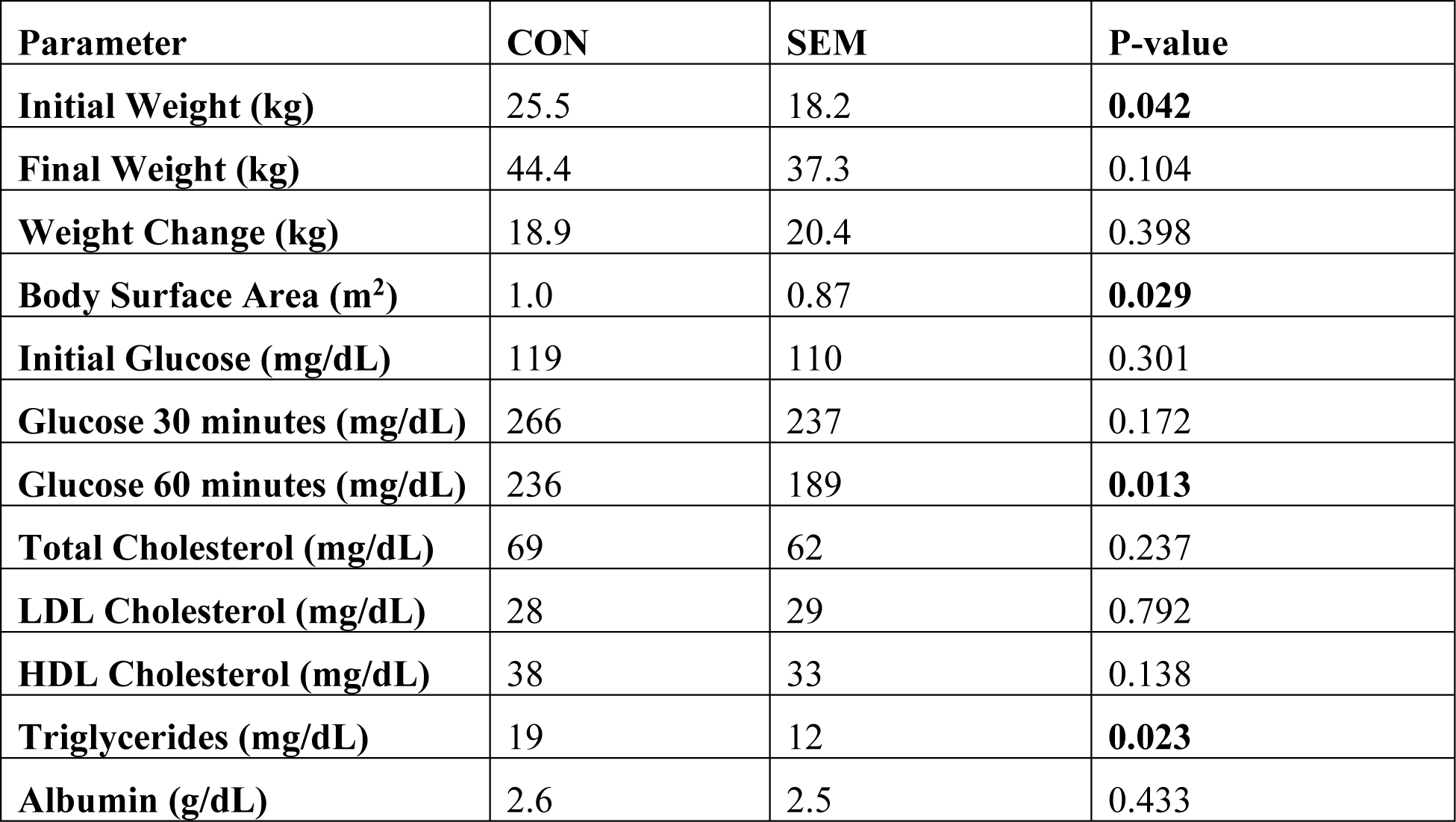
Physical and Metabolic Parameters. Semaglutide-treated animals (SEM) exhibited improved glucose tolerance and serum triglycerides compared to controls (CON), and no difference with respect to cholesterol, albumin, or body weight. Body surface area was significantly reduced in SEM animals at the end of our experimental protocol. Statistically significant values are bolded.

**Table 2.**
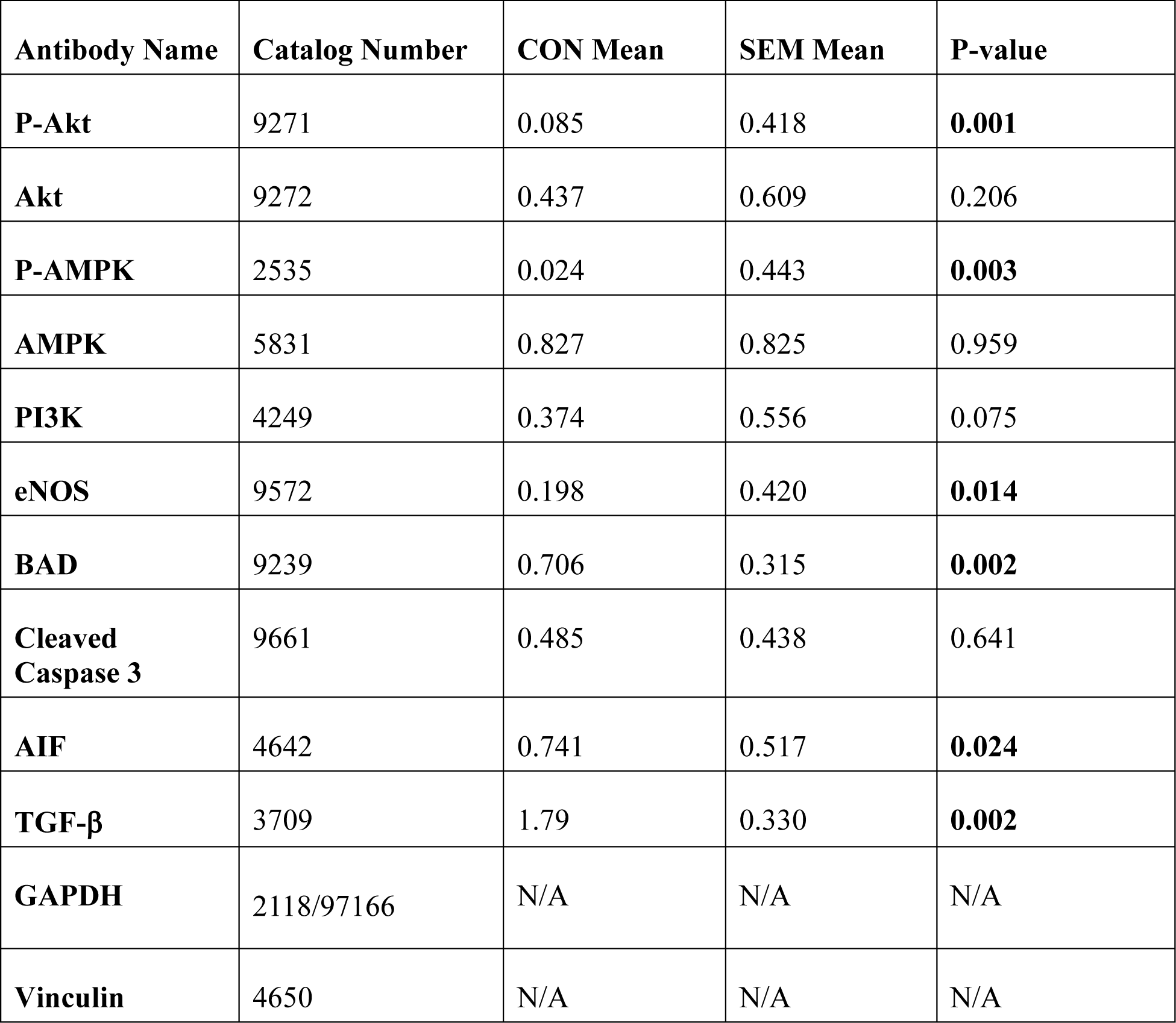
Antibodies utilized in this experiment. All antibodies are sourced from Cell Signaling (Danvers, MA). P-values were calculated following normalization for protein loading. See Materials and Methods for additional detail regarding immunoblotting and statistical testing, and Figure S1 for the band images that were analyzed to yield the results described in this table. P-Akt: phospho-AKT, Ser473. P-AMPK: phospho-AMP-activated protein kinase-α, Thr172. PI3K: phosphatidylinositol-3-kinase, p110α. eNOS: endothelial nitric oxide synthase. BAD: bcl- 2-associated death promoter. AIF: apoptosis-inducing factor. TGF-β: transforming growth factor-β. GAPDH: glyceraldehyde-3-phosphate dehydrogenase. P-values that met the predetermined threshold for significance are bolded.

### Semaglutide Improves Systolic Function in the Chronically Ischemic Heart

Pressure- volume (PV) loop values derived from direct left ventricular chamber cannulation revealed significant improvements in left ventricular ejection fraction in SEM animals, both at rest and during rapid ventricular pacing, when compared with CON values (40.4 vs. 27.0% left ventricular volume, p=0.025 at rest, and 30.1 vs. 13.0% left ventricular volume during rapid pacing). This augmentation in ejection was accompanied by a numerically increased cardiac index, both at rest and paced (3.48 vs. 2.09 L/min/m^2^, p=0.051 at rest, and 4.77 vs. 2.18 L/min/m^2^, p=0.103 during pacing), that trended toward but did not reach the pre-specified threshold for statistical significance set at p<0.05. Similarly, diastolic function as delineated by Tau, or the isovolumetric ventricular relaxation constant, was numerically improved in SEM animals, but not statistically so (24.3 ms vs. 30.4 ms, p=0.061). Incorporation of inferior vena cava occlusion-based PV loop measurements permitted acquisition of multiple load-independent measures of intrinsic cardiac inotropy, including the end-systolic pressure-volume relationship (ESPVR) and the preload-recruitable stroke work slope (PRSW); neither the ESPVR nor the PRSW was altered in the SEM group compared to CON (3.04 vs. 2.46, p=0.497, and 40.9 vs. 43.7, p=0.645, respectively). Finally, there was no difference in heart rate or in mean arterial pressure between treatment groups (p=0.418 and p=0.460, respectively). Figure 1 depicts these changes visually.

**Fig. 1.**
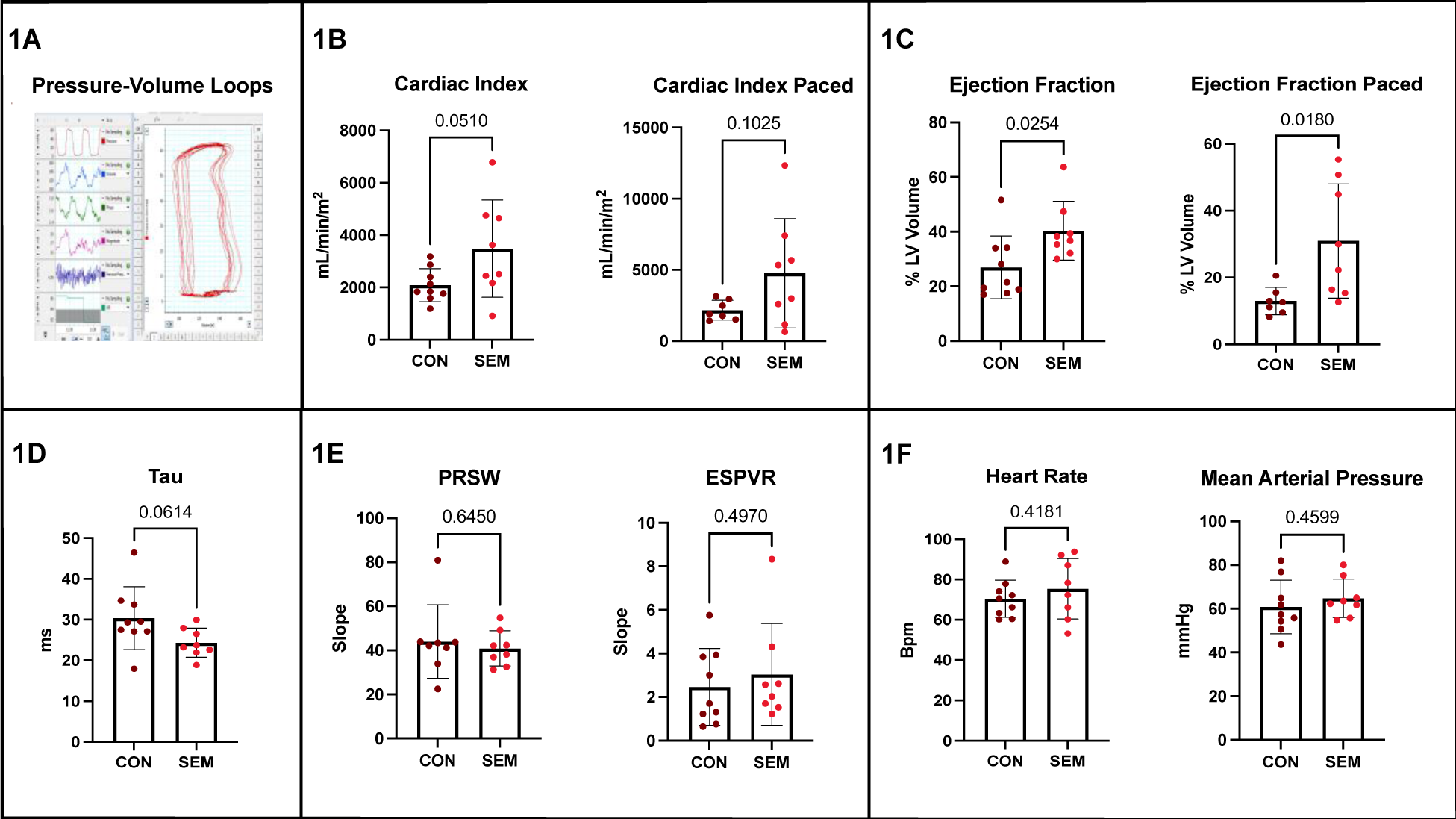
Pressure-volume loop-acquired hemodynamic measurements. **1A:** Example intraoperative pressure-volume loops captured from the transduction software display. These tracings generate the chamber parameters ultimately productive of a comprehensive picture of cardiac function and, through loop morphology, immediate feedback regarding the propriety of catheter positioning. **1B:** Semaglutide-treated animals (SEM) exhibited improved left ventricular ejection fraction, both at rest and under simulated cardiac stress using rapid pacing, when compared with control animals (CON). **1C:** There were trends toward improved cardiac index at rest and with pacing. **1D:** This was also true of diastolic function as indicated by Tau, the isovolumetric ventricular relaxation constant. **1E:** No changes were observed in intrinsic contractility as assessed by the end-systolic pressure-volume relationship (ESPVR) or by the preload-recruitable stroke work slope (PRSW). **1F:** There was no difference in baseline heart rate or mean arterial pressure between groups. Each dot represents an individual animal.

### Semaglutide Promotes Perfusion of the Ischemic Myocardium

Left ventricular segments determined to represent the most ischemic cardiac foci using neutron activation assays for gold microsphere quantity were subsequently analyzed for perfusion. Neutron activation assays were also conducted for additional microsphere isotope quantities, with unique isotopes injected at rest and during rapid myocardial pacing to assess differential perfusion under these conditions; see the Materials and Methods section for the mathematical derivation of perfusion from isotopic quantity. Using this method, SEM animals exhibited significantly elevated coronary perfusion to the territory rendered most ischemic by ameroid constrictor placement at resting heart rate (0.89 vs. 0.41 mL/min/g, p=0.017), and this elevation was maintained in the context of myocardial stress by rapid pacing (0.95 vs. 0.35 mL/min/g, p=0.026). Because GLP-1 agonist-associated myocardial protection has been linked to AMPK pathway activation and nitric oxide-mediated vasodilation, associated markers were probed by immunoblotting on tissue from the most ischemic myocardial segments (13, 30). This revealed significant increases in activated AMPK expression (p=0.003), activated AKT expression (p=0.001), and endothelial nitric oxide synthase expression (p=0.014), implicating augmented vasodilatory efficacy as an account of the observed improvements in perfusion. Figure 2 depicts perfusion graphically in conjunction with associated immunoblot images. Antibody details, along with additional immunoblot images and values, including the GAPDH or vinculin probed on each membrane to control for loading error, may be found in Supplementary Figure 1 and Table 2. Additionally, Movie 1 provides an intraoperative depiction of the procedural derivation of these data.

**Fig. 2.**
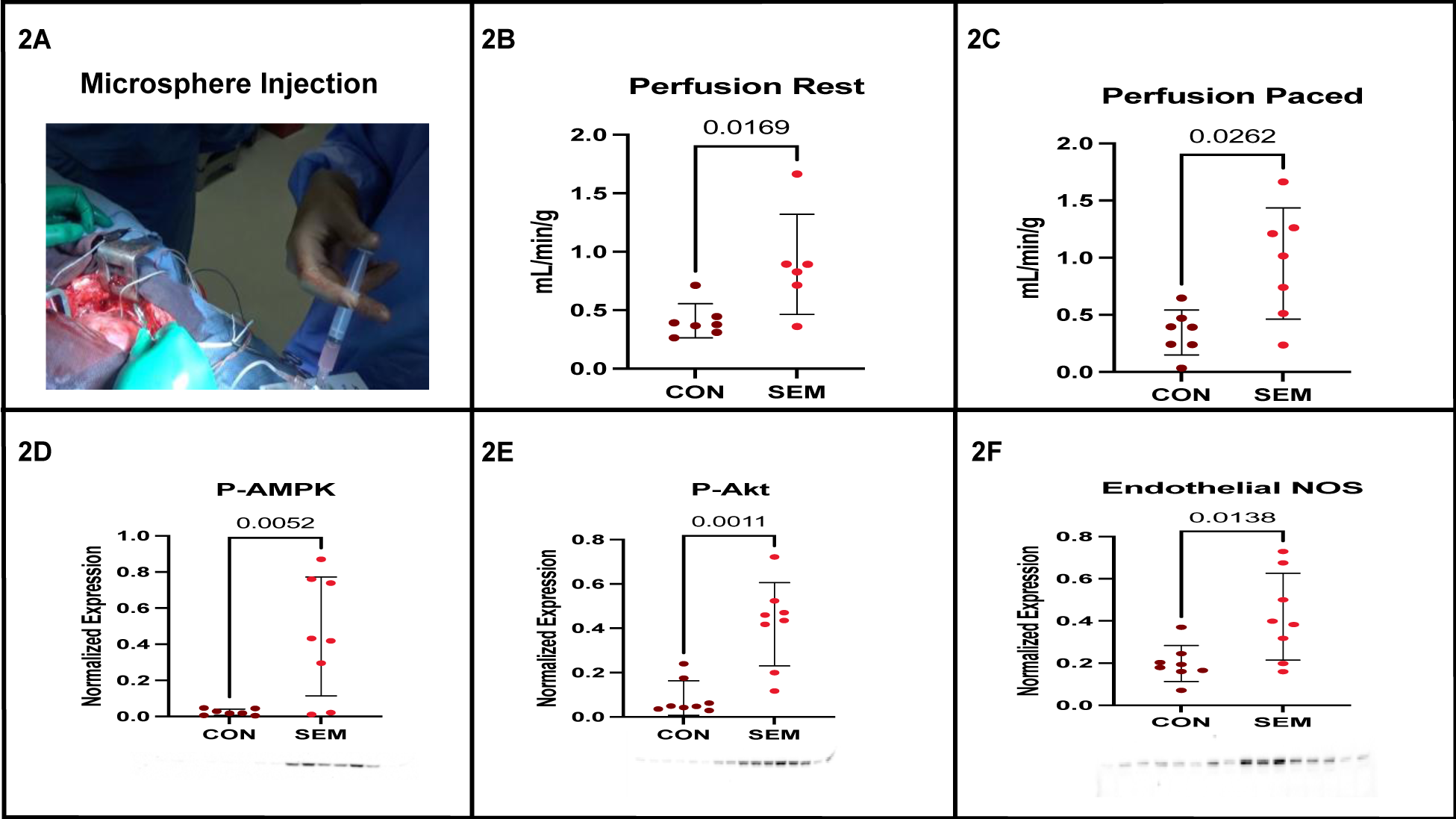
Perfusion differences and associated molecular markers. **2A:** Intraoperative photograph depicting isotopically labelled microsphere injection into the left atrial appendage. Unique isotopes are injected at resting heart rate and during rapid myocardial pacing; subsequent separate analysis following neutron activation permits derivation of perfusion during both conditions. **2B:** Perfusion to the most ischemic myocardial territory assessed by these means revealed significant augmentation in favor of semaglutide-treated animals (SEM), in which group perfusion was increased 2.2-fold over control (CON) equivalents. **2C:** Perfusion to the most ischemic myocardial territory during rapid pacing was also significantly increased in SEM animals, in this case representing a 2.7-fold increase when compared with the CON mean. **2D:** Immunoblotting revealed a marked increase in phospho-AMPK (p-AMPK) expression; the analyzed band image appears below, with bands located below the treatment group from which they originated. **2E:** There was also a significant increase in phospho-Akt (p-AKT) expression favoring the SEM group; the analyzed band image appears below, with bands located below the treatment group from which they originated. **2F:** Endothelial nitric oxide synthase (NOS) expression was significantly increased in SEM animals as well; the analyzed band image appears below, with bands located below the treatment group from which they originated, with dots representing unique lysates. All immunoblots are reported as relative expression, as all expression was normalized to GAPDH or vinculin probed on the same membrane.

### Semaglutide Supports Cellular Survival in the Ischemic Myocardium

Reduced apoptosis in general and BAD inhibition in particular has also been implicated in the cardioprotective effects of GLP-1 agonist therapy. We employed immunofluorescent TUNEL staining and immunoblotting for apoptotic markers within the most ischemic myocardial sections of all animals (14, 25). Compared with the CON mean, there was a marked reduction in apoptotic fragments by TUNEL immunofluorescence in SEM myocardial sections (p=0.008). In corroboration of the mechanism established elsewhere in the literature, this was accompanied on immunoblot analysis by significant expression reductions in the pro-apoptotic markers BAD and apoptosis-inducing factor (AIF, p=0.002 and p=0.024, respectively). There was no significant difference in cleaved caspase 3 expression (p=0.641). Figure 3 displays representative TUNEL immunofluorescence-subjected slides from both groups, along with associated immunoblots.

**Fig. 3.**
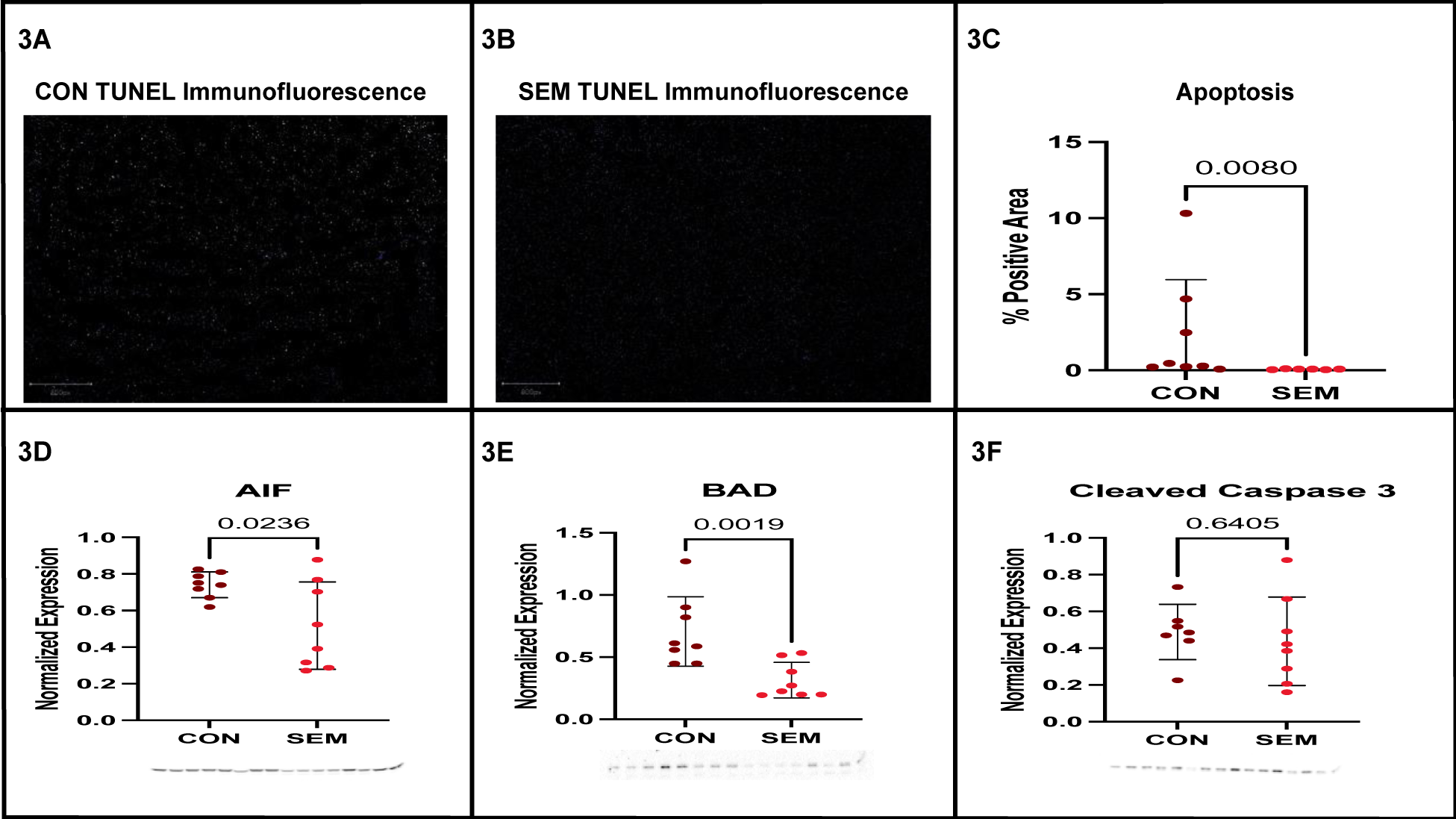
Myocardial apoptotic burden according to treatment group. Terminal deoxynucleotidyl transferase dUTP nick-end labeling (TUNEL) with fluorescein was applied to the most ischemic myocardial sections of all animals to compare the apoptotic burden according to treatment group; images were analyzed by independent technicians blinded to experimental circumstances. **3A:** A representative image depicting apoptotic cell death within control group (CON) tissue. **3B:** A representative image depicting apoptosis within semaglutide-treated (SEM) tissue. **3C:** Analysis of percent area affected by apoptosis averaged across sections from all animals revealed significantly reduced apoptosis in the SEM group. **3D:** Immunoblotting demonstrated significantly attenuated apoptosis-inducing factor (AIF) expression in SEM tissue when compared with the CON mean. **3E:** There was also a significant reduction in bcl-2- associated death promoter (BAD) expression in SEM tissue when compared with the CON mean. **3F:** There was no difference in cleaved caspase 3 expression between the SEM and CON groups. All immunoblots are reported as relative expression, as all expression was normalized to GAPDH or vinculin probed on the same membrane. The scale bar in the lower left corner of both immunofluorescent images represents 800 pixels.

### Semaglutide Attenuates Interstitial and Perivascular Fibrosis in the Ischemic Myocardium

Given the augmented perfusion found in semaglutide-treated ischemic myocardial tissue sections in conjunction with the cardiac anti-fibrotic effect attributed to GLP-1 receptor agonism in the literature, a reduction of the fibrotic burden in this tissue would not be unanticipated ^39^. When subjected to Masson’s trichrome protocol to quantify collagenous deposits within the ischemic myocardial interstitium and surrounding its arteriolar supply, we observed significant reductions in both locations in the SEM group when contrasted with CON controls (9.9 vs. 2.9% positive area, p=0.020, and 49 vs. 9.1% positive area, p=0.002, respectively). In concordance with this multifocal alleviation of collagen deposition and with the mechanism reported in the literature, probing the ischemic myocardial tissue for TGF-β protein expression revealed a significant reduction in SEM animals (p=0.002). Figure 4 depicts these differences, their quantification by area, and associated immunoblotting data.

**Fig. 4.**
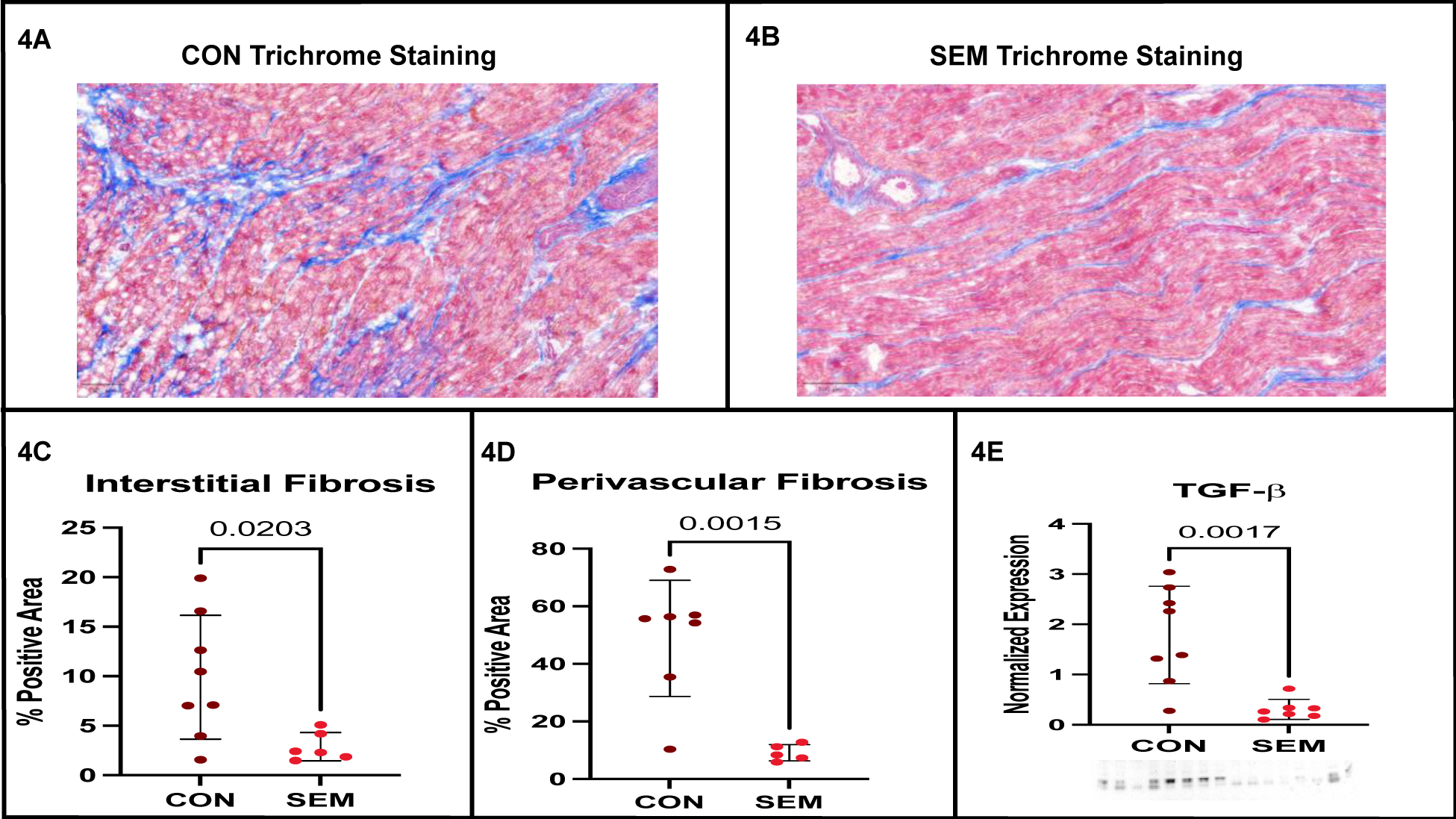
Myocardial interstitial and perivascular fibrosis according to treatment group. Masson’s trichrome staining was used to render comparative fibrotic burden within the myocardial interstitium and its arteriolar supply in sections taken from the most ischemic left ventricular segments of semaglutide-treated (SEM) and control (CON) animals. Collagen fibers appear in blue, cardiomyocytes in red, and vascular smooth muscle in magenta. The bar in the lower left corner of both histologic images represents 100 μm. **4A:** Trichrome staining in the most ischemic myocardial section taken from a CON animal. **4B:** Trichrome staining in the most ischemic myocardial section taken from a SEM animal. **4C:** There was a significant (3.4-fold) reduction in interstitial fibrosis in SEM animals. **4D:** There was also a significant (5.4-fold) reduction in perivascular fibrosis in SEM animals. **4E:** This multifocal reduction in fibrotic burden occurred in association with a significant reduction in TGF-β expression in SEM animals.

## Discussion

With statements such as the *New Yorker*’s “Year of Ozempic” appellation for the year 2023, there are few medical phenomena in recent memory that have captured the cultural moment as completely as these drugs (32, 33). So far, much of the focus of the field has centered around, rather than the original impetus underlying the development of incretin-based agents of achieving glycemic control, the closely associated and pleiotropic metabolic benefits of these drugs. Beginning with FDA approval of liraglutide for chronic weight management nearly ten years ago, countless trials have since documented a spectrum of associated benefits, including in conditions as diverse as nonalcoholic steatohepatitis, chronic kidney disease, obesity-related heart failure, and even Parkinson’s disease ^42–45^.

Of all manners of metabolic betterment attributed to these drugs, however, there is none poised to impose as profound an impact as that affecting cardiovascular disease. Precise accounting of the magnitude of this effect is precluded by the recency of GLP-1 agonist approval for cardiovascular risk reduction in March of 2024, but meta-analytic evidence ranging over 60,000 diabetic patients and demonstrating a 15% reduction in major adverse cardiac events, a 16% reduction in myocardial infarction, a 12% reduction in cardiovascular death, and a 17% reduction in all-cause mortality, indicates strong potential for improvement in quality and quantity of life for millions of patients with cardiovascular disease ^9^.

The publication of a recent major trial in non-diabetic obese patients with pre-existing cardiovascular disease demonstrating that the cardiovascular risk reduction seen with use of GLP-1 agonist drugs appears to arise independently of effects on glycemic control, however, raised an additional question that has not yet been adequately addressed ^10^: are the cardiovascular benefits of these drugs due to systemic risk factor mitigation, or do they truly represent direct, disease-modifying effects on cardiovascular tissue? This is not a merely academic question. While all of the large trials that studied GLP-1 agonists and cardiovascular disease risk were conducted in patients with either diabetes or obesity, only 29% of patients with CAD have diabetes, while only 33% of patients with CAD are also obese ^46,47^. In other words, the prospect that these drugs may benefit all patients with heart disease, and not just those with certain comorbid conditions, remains clinically unexplored.

In our experiment we sought, by employing an extensively validated large animal model ^35^ devoid of potentially confounding metabolic comorbidities, to supply evidence germane to the assessment of this prospect in the context of chronic coronary artery disease. The consequence of the ameroid constrictor placed around the left circumflex coronary artery during the initiating surgery of this model is the gradual induction, over the course of several weeks, of ischemic cardiomyopathy affecting approximately 20% of the left ventricular muscle if placed, as in this experiment, as proximally as possible to the arterial origin ^48^. This proportion of myocardium is thereby rendered, given the relative paucity of epicardial coronary collaterals that swine share with humans, hypoperfused, dyskinetic, and ultimately infarcted, in proportion to the absence of circulatory compensation. In our study, perfusion to the most ischemic myocardial territory and systolic cardiac function (as indicated by left ventricular ejection fraction) were significantly augmented together in animals that received semaglutide (SEM) compared with counterparts that did not (CON). These concurrent findings indicate the production of a more robust myocardial compensatory response with GLP-1 agonist treatment. The collateral circulation that develops in response to ameroid-mediated ischemia has been found to lack the arteriolar smooth muscle and vasodilatory response needed to match coronary flow to exercise-induced stress, replicating the pathology responsible for impaired exercise tolerance in patients with coronary ischemia ^32^. The additional finding in our study that semaglutide-associated augmentation persisted during the subjection of hearts to rapid ventricular pacing is important in this regard, as it suggests the cultivation of a circulatory reserve capacity capable of interrupting the induction of exercise-induced angina in patients ^49^.

An additional component of our results relevant to their translational value is the equivalence of multiple sensitive, load-independent indices of myocardial contractility between the SEM and CON animals. This is consistent with previous preclinical experimentation, and implies that the improvement in myocardial performance seen with semaglutide is mediated by Frank-Starling dynamics that do not emerge at the energetic expense of a cardiomyopathic heart ^50^. The trend towards reduced Tau seen in SEM animals, which is a measure of ventricular relaxation and diastolic function, supports this contention ^51^. Finally, the unchanged heart rate and mean arterial pressure between groups further demonstrates that the favorable functional effects of GLP-1 agonist therapy do not require toleration of adverse implications on afterload or myocardial work. These are crucial considerations in contemplating the application of these findings to patients with ischemic heart disease, in many of whom an incidental hemodynamic decrement would preclude trial of an additional drug ^52^. In other words, because semaglutide does not appear to produce functional improvement at the expense of additional myocardial oxygen demand, its incorporation would not compromise the purpose of the current therapeutic paradigm, which aims to balance myocardial energetic expenditure with the impaired supply of nutrients imposed by flow-limiting ischemia ^53^.

The histologic and molecular data generated during our experiment both corroborate previous accounts from the literature of the cardioprotective effects of GLP-1 agonist administration, and elucidate our novel findings mechanistically. There are many studies documenting the endothelial-protective effects of GLP-1 agonists that, taken together, suggest that upregulation of the Akt-AMPK-eNOS pathway produces, through increased nitric oxide elaboration, improved coronary blood flow. This, in turn, reduces the myocardial ischemic burden and bolsters cardiac function ^54^. In our study, upregulation of this pathway in a large animal model with clinically relevant cardiac anatomic and physiologic homology to the human heart represents a substantial extension of the applicability of this account to patients, as it has been previously shown only in small animals, cell culture, and in human peripheral vasculature ^55^.

As with the molecular underpinnings of the perfusion changes found to arise in the SEM cohort, the improved myocardial cellular survival and blunted fibrotic remodeling we observed provide simultaneous corroboration and extension of the established account of GLP-1 agonism in the heart. Reductions in apoptotic cell death have been found with GLP-1 agonist administration acutely in an ischemia-reperfusion model, but our study is the first to demonstrate the longevity of this reduction in the setting of a model of chronic myocardial ischemia reminiscent of the disease process as it transpires in patients ^14^. This is true also of the marked reductions in perivascular and interstitial myocardial fibrosis we observed in SEM animals: despite a profusion of preclinical studies in small animals and cells demonstrating that GLP-1 agonists interrupt fibrotic pathophysiology and reduce remodeling in the wake of injury, these findings have never been reported following a clinically meaningful duration of disease ^39^. Given this, the functional improvements produced by semaglutide in our study may involve mitigation of the fibrotic disruption of the myocardial contractile apparatus that occurs after ischemic injury ^56^.

Metabolic and physical parameters collected over the course of the experiment suggest that salubrious systemic changes may have operated in parallel with these direct intramyocardial effects to culminate in the functional improvements recorded in our study. Although there was no difference in weight or in weight change between CON and SEM animals, the significantly reduced triglycerides seen in SEM animals has both been found in patients undergoing GLP-1 agonist therapy, and represents the reduction of a known cardiovascular risk factor ^37^. This, along with the improved glucose tolerance seen in the SEM group despite the absence of baseline hyperlipidemia or hyperglycemia in these animals, underscores the potential of GLP-1 agonist therapy to serve as a protective factor against the development of these known metabolic risks associated with incident CAD and worsening of the phenotype thereof ^57^; this prospect, while of interest to the specification of indications for GLP-1 agonist therapy in patients with cardiovascular disease, requires longer-term clinical follow-up for validation.

In summary, even when assessed in a model structured by induction of chronic myocardial ischemia in the absence of potentially confounding comorbidities, oral semaglutide delivered in a clinically appropriate dose and duration produced significant improvements in coronary perfusion, cardiac performance, myocardial cellular survival, and tissue architecture. These findings demonstrate the directly cardioprotective efficacy of GLP-1 agonist therapy, and warrant consideration of the use thereof in patients with ischemic heart disease, regardless of the presence of concomitant metabolic conditions.

## Non-standard Abbreviations and Acronyms

CAD: coronary artery disease
GLP-1: glucagon-like peptide 1
eNOS: endothelial nitric oxide synthase
LCx: left circumflex coronary artery
TUNEL: terminal deoxynucleotidyl transferase dUTP nick-end labeling
AMPK: 5’-adenosine monophosphate-activated protein kinase
PI3K: phosphatidylinositol-3-kinase, p110α
BAD: bcl-2-associated death promoter
AIF: apoptosis-inducing factor
TGF-β: transforming growth factor-β
BCA: bicinchoninic acid assay
GAPDH: glyceraldehyde-3-phosphate dehydrogenase
ESPVR: end-systolic pressure-volume relationship
PRSW: the preload-recruitable stroke work slope

## Acknowledgements

The authors would like to extend their deep gratitude to the veterinary staff of Brown University and Rhode Island Hospital for their indispensable physiological erudition, dedicated animal husbandry, and perioperative assistance. Without the steadfast support of this consummately professional team, the experimental and ethical standards achieved in this work would have been unattainable. Sources of funding: National Institutes of Health grant T32HL160517 (CRS, DDH, MB); Armand D. Versaci Research Scholar in the Surgical Sciences Award (MK); National Institutes of Health grant R01HL133624 (MRA); National Institutes of Health grant R56HL133624-05 (MRA); National Institutes of Health grant R01HL46716 (FWS); National Institutes of Health grant R01HL128831 (FWS); Rhode Island Foundation grant 1472420231352 (MRA). Disclosures: none.

## Supplemental Material

Figure S1

Movie 1

## Highlights

– When administered in the setting of a clinically relevant large animal model of chronic myocardial ischemia, semaglutide improved myocardial systolic performance and perfusion to the ischemic myocardial territory
– The improved perfusion observed in our study appeared to arise mechanistically from Akt-AMPK-eNOS pathway regulation, and was also associated with reduced perivascular fibrosis and myocardial cellular survival
– The occurrence of these favorable changes in the context of a model of coronary artery disease alone, in the absence of the cardiometabolic comorbidities to which semaglutide use is currently confined, suggest that the applicability of this drug may be wider than is currently conceived

## Notes

### Competing Interest Statement

The authors have declared no competing interest.

